# Kaposi’s sarcoma-associated herpesvirus ORF66 is essential for late gene expression and virus production via interaction with ORF34

**DOI:** 10.1101/728147

**Authors:** Tadashi Watanabe, Mayu Nishimura, Taisuke Izumi, Kazushi Kuriyama, Yuki Iwaisako, Kouhei Hosokawa, Akifumi Takaori-Kondo, Masahiro Fujimuro

## Abstract

Kaposi’s sarcoma-associated herpesvirus (KSHV) is closely associated with B-cell and endothelial cell malignancies. After the initial infection, KSHV retains its viral genome in the nucleus of the host cell and establishes a lifelong latency. During lytic infection, KSHV encoded lytic-related proteins are expressed in a sequential manner and are classified as immediate early, early, and late gene transcripts. The transcriptional initiation of KSHV late genes is thought to require the complex formation of the virus specific pre-initiation complex (vPIC), which may consist of at least 6 transcription factors (ORF18, 24, 30, 31, 34, and 66). However, the functional role of ORF66 in vPIC during KSHV replication remains largely unclear. Here, we generated ORF66-deficient KSHV using a BAC system to evaluate its role during viral replication. While ORF66-deficient KSHV demonstrated mainly attenuated late gene expression and decreased viral production, viral DNA replication was unaffected. CHIP analysis showed that ORF66 bound to the promoters of late gene (*K8.1*), but did not to those of latent gene (*ORF72*), immediate early gene (*ORF16*) and early gene (*ORF46/47*). Furthermore, we found that three highly conserved C-X-X-C sequences and a conserved leucine-repeat in the C-terminal region of ORF66 were essential for interaction with ORF34 and viral production. The interaction between ORF66 and ORF34 occurred in a zinc-dependent manner. Our data support a model, in which ORF66 serves as a critical vPIC component to promote late viral gene expression and viral production.

**IMPORTANCE:** KSHV ORF66, a late gene product, and vPIC are thought to contribute significantly to late gene expression during the lytic replication. However, the physiological importance of ORF66 in terms of viral replication and vPIC formation remains poorly understood. Therfore, we generated a ORF66-deficient BAC clone and evaluated its viral replication. Results showed that ORF66 played a critical role in virus production and the transcription of L genes. To our knowledge, this is the first report showing ORF66 function in virus replication using ORF66-deficient KSHV. We also clarified that ORF66 interacted with the transcription start site of *K8.1* gene, a late gene. Furthermore, we identified the ORF34-binding motifs in the ORF66 C-terminus: three C-X-X-C sequences and a leucine-repeat sequence, which are highly conserved among β- and γ-herpesviruses. Our study provides insights into the regulatory mechanisms of not only the late gene expression of KSHV but also those of other herpesviruses.

## Introduction

KSHV, (also known as human herpesvirus 8 or HHV-8), causes Kaposi’s sarcoma, primary effusion lymphoma (PEL), and multicentric Castleman’s disease (1–3). Since the first discovery of KSHV DNA fragments in a Kaposi’s sarcoma lesion of an AIDS patient in 1994 (4), over a quarter of a century has passed, and several aspects of KSHV pathogenesis, life-cycle and viral protein function have been elucidated. Compared with other viruses, herpesvirus has a large number of viral genes in its genome. KSHV encodes not only viral proteins but also miRNAs and LncRNAs. These viral molecules are thought to be essential for KSHV replication and its pathogenesis. One characteristic of the KSHV life-cycle is the establishment of a lifelong latency in the infected individual leading to KSHV-associated malignancy in patients with severe immunosuppression by drugs after organ transplantation or AIDS (1–5). Development of KSHV-associated neoplasm occurs due to infected cells which express few latent KSHV genes (including LANA, v-FLIP, Kaposin, and miRNAs) (6). These genes modulate cell-proliferation and apoptosis pathways (7–11). Another characteristic of the KSHV life-cycle is active virus production, known as lytic infection. The genes related to lytic infection have been categorized into into three groups, immediate early (IE), early (E), and late (L) (12, 13). Sequential and temporal expression of KSHV genes are key to induce efficient viral production during lytic infection. The IE-gene product RTA/ORF50 is a transcription factor for triggering transcriptional activation of another IE genes and E genes, and RTA/ORF50 initiates the shift from latency to lytic infection (13, 14). The transcribed products of E genes start viral genome DNA replication. Finally, the transcriptional products of L genes, contribute to viral particle formation by its encoding viral structure proteins (15).

The viral pre-initiation complex (vPIC) has recently been proposed to regulate L gene expression(16). vPIC is composed of several viral proteins conserved among β- and γ-herpesviruses (17), and has functional homology with the host pre-initiation complex, which consists of TATA-binding protein (TBP) and general transcriptional factors (GTFs). Whereas the host pre-initiation complex accumulates on the TATA-box of transcriptional start site (TSS) and initiates cellular RNA polymerase II (RNA pol II)-mediated transcription, vPIC accumulates on the “TATT”-box of viral gene TSS and initiates RNA pol II-mediated transcription (17).

The function of vPIC machinery and its components has been extensively studied in the context of EBV, MHV-68, CMV and KSHV(16). In KSHV, at least 6 viral proteins contribute to vPIC formation. Viral TBP homolog KSHV ORF24 directly binds to the “TATT”-box on the promoter sequences of the KSHV genome and is essential for the recruitment of host RNA pol II (18, 19). We and other groups revealed that KSHV ORF34 acts as a hub for interaction between ORF24 and other vPIC components such as ORF18, ORF30, ORF31, and ORF66 (18, 20, 21). Split luciferase assays and co-immunoprecipitation experiments revealed that ORF34 directly/or indirectly interacts with ORF 18, 30, 31 and 66 (18, 20–22). ORF24 binds to the promoter of the L genes with RNA pol II, and ORF34 serves as a bridge between ORF24 and a complex of ORF18, 30, 31 and 66 (18, 20, 21). Furthermore, ORF18 (23), ORF30 (21), ORF31 (23) and ORF34 (20) are essential for virus replication and gene expression. Although ORF66 appears to be a vPIC component (18, 20, 22), the importance and its function during KSHV replication remains unknown. Therefore, we established ORF66-deficient KSHV, and evaluated its physiological role during viral replication. Here, we show that ORF66 is essential for virus production and L gene expression via interaction with ORF34.

## Results

### Construction of ORF66-deficient KSHV BAC

We constructed ORF66-deficient recombinant KSHV BAC (ΔORF66-BAC16) to study the impact of ORF66 during KSHV replication. Three stop codons (3-stop element) were inserted into the ORF66 coding region of KSHV BAC16 using a two-step markerless red recombination system (24, 25) (Fig. 1a). Because ORF66 overlaps with ORF67 (Fig. 1a), the 3-stop codons were inserted within the ORF66 gene to avoid interference with the coding frame of ORF67. The insertion and deletion of a kanamycin resistance gene (Kan^R^) were analyzed by EcoRV-digestion (Fig. 1b). Mutations and the insertion of the 3-stop codons into ΔORF66-BAC16 were confirmed by Sanger sequencing (Fig. 1c).

**Fig.1.**
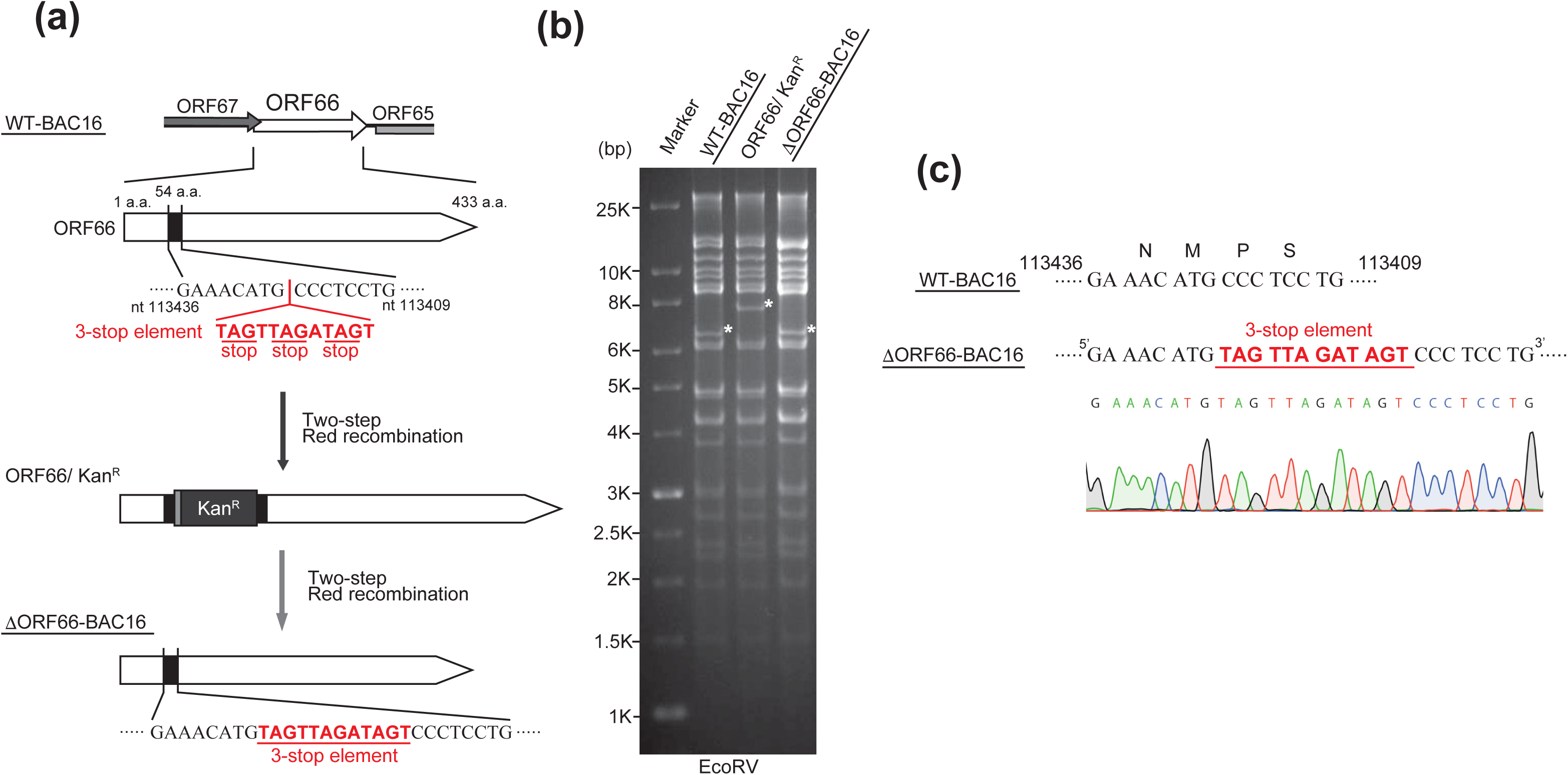
Construction of recombinant ΔORF66 KSHV BAC. (a) Schematic illustration of the KSHV genome including the ORF66 coding region. Using a two-step Red recombination system, three stop codons were inserted into the ORF66 coding region of KSHV BAC16 (nt113417 - nt113416; Accession number: GQ994935) to construct ORF66-deficient BAC clone (ΔORF66-BAC16). (b) Agarose gel electrophoresis of the recombinant KSHV BACmids, digested with EcoRV. The asterisks (*) indicate insertion and deletion of a kanamycin-resistance cassette in each BAC clone. (c) DNA sequencing results of ORF66 mutagenesis sites in ΔORF66-BAC16.

### ORF66 deficiency abrogates virus production and late gene expression

To efficiently induce recombinant KSHV, tetracycline-inducible (Tet-on) RTA/ORF50-expressing SLK cells (iSLK) and Vero cells (iVero) were used as virus producer cells (20). Recombinant KSHV BAC clone, wild type (WT)-BAC16 or ΔORF66-BAC16, was transfected into iVero or iSLK cells, and then selected with hygromycin to generate recombinant KSHV-inducible stable cell lines, iVero-WT, iVero-ΔORF66, iSLK-WT, and iSLK-ΔORF66. To evaluate whether ORF66 is critical for KSHV replication, virus production and virus genome replication in iSLK-ΔORF66 and iVero-ΔORF66 cells were analyzed. iSLK-WT, iSLK-ΔORF66, iVero-WT, and iVero-ΔORF66 cells were treated with Dox and NaB, and culture supernatants were harvested. The amount of WT-KSHV and ΔORF66-KSHV in iSLK or iVero culture supernatants were measured by real-time PCR (Fig. 2a and 2e). As a result, the production of ΔORF66-KSHV in iSLK and iVero cells were about 1000-fold and 100-fold lower than WT-KSHV. In contrast, there was no significant difference in cell-associated KSHV DNA levels between WT-KSHV and ΔORF66-KSHV producing cells (Fig. 2b and 2f). Next, to clarify that the reduction of virus production in iSLK-ΔORF66 and iVero-ΔORF66 cells is caused by ORF66-deficiency, we tested whether exogenous ORF66 expression could rescue virus production of iSLK- and iVero-ΔORF66 cells. The iSLK-ΔORF66 and iVero-ΔORF66 cells stably transfected with empty plasmid or ORF66 expression plasmid were cloned by neomycin (G418) or blastcidin S. The protein expression levels of exogenous 3xFLAG-tagged ORF66 were confirmed by Western blotting (Fig. 2c and 2g). Virus production from iSLK-ΔORF66 and iVero-ΔORF66 cell lines was partially but significantly recovered when 3xFLAG-ORF66 was exogenously expressed (Fig.2d and 2h). These results indicate that ORF66 is crucial for KSHV replication and may function in steps following viral DNA replication.

**Fig.2.**
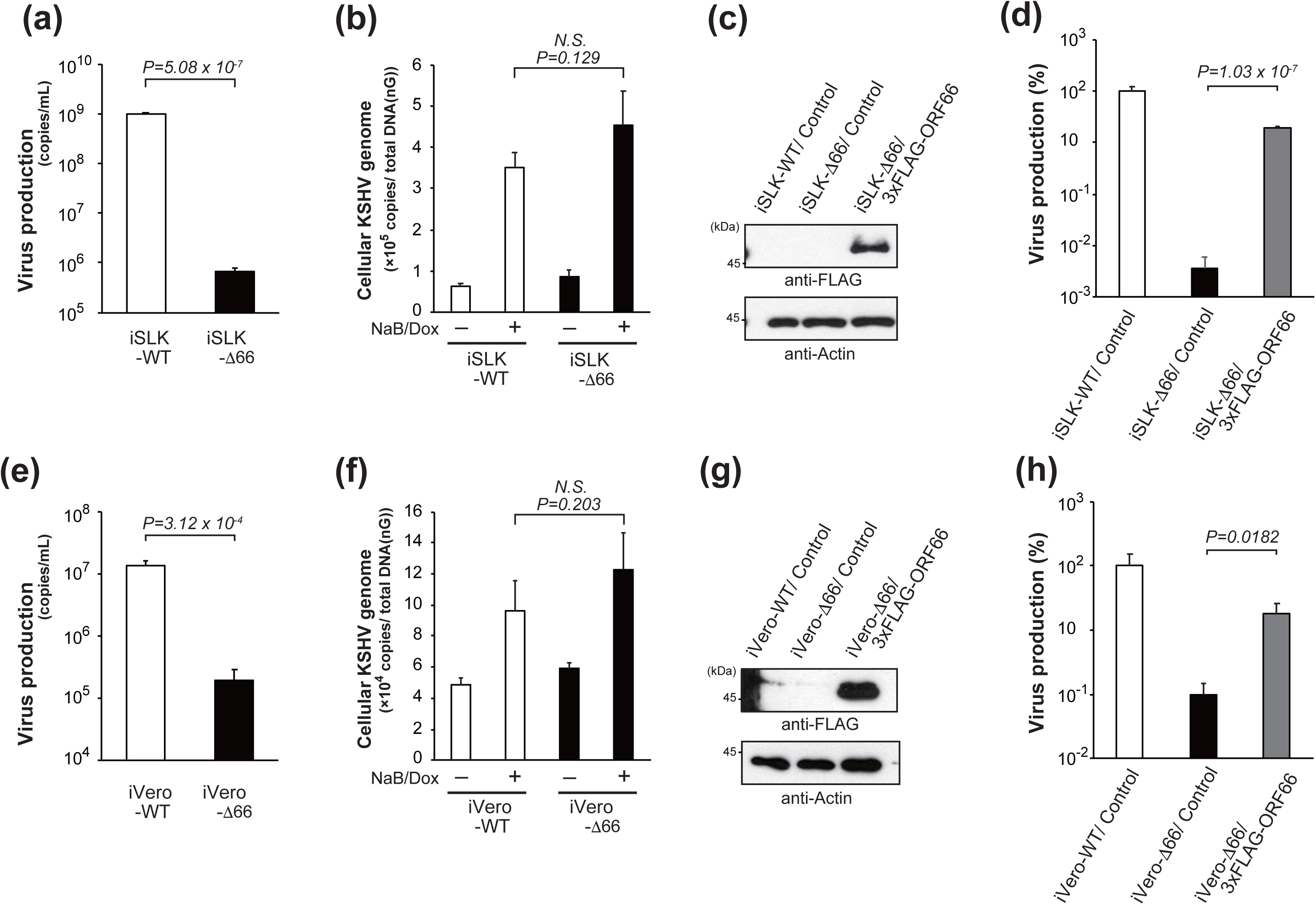
ORF66 is essential for virus production but not DNA replication of KSHV. Virus production in (a) iSLK-WT, iSLK-ΔORF66 and (e) iVero-WT, iVero-ΔORF66. Each cell line was cultured for 72 h with medium containing NaB and Dox. KSHV DNA was purified from capsidated KSHV virions in culture supernatants, and KSHV genome copies were determined by real-time PCR. Virus production in (b) iSLK-WT, iSLK-ΔORF66 and (f) iVero-WT, iVero-ΔORF66. Each cell line was cultured for 48 h with medium containing of NaB and Dox. Cellular DNA containing KSHV genomic DNA was purified from each cell line. KSHV genome copies were determined by real-time PCR and normalized by the total DNA amount. Establishment of exogenous ORF66 expressing (c) iSLK-ΔORF66 and (h) iVero-ΔORF66 stable cell lines. Western blot shows exogenous FLAG-tagged ORF66. Rescue of virus production in (d) iSLK-ΔORF66 and (h) iVero-ΔORF66 cells by exogenous ORF66 expression. Each stable cell line was cultured with NaB and Dox-containing medium for 3 days, and culture supernatant containing virus was harvested and quantified. (a-b, d-f, h) Three or four independent samples were evaluated by real-time PCR. The error bars indicate standard deviations.

To evaluate the contribution of ORF66 on KSHV gene expression, we performed an RT-qPCR array on viral gene. Total RNA was extracted from iSLK-WT and iSLK-ΔORF66, stimulated with NaB and Dox for 72 hours. RNA was subjected to RT-qPCR array as previously reported (26). Our data showed a broad reduction of KSHV mRNA in iSLK-ΔORF66 compared to iSLK-WT (Fig.3), where 7 out of the top 10 down-regulated genes were late genes (*K8.1*, *ORF17*, *ORF26*, *ORF27*, *K9*, *ORF75*, *ORF25*). In particular, *K8.1*, *ORF26*, *ORF27, and ORF25* have been previously reported by Nandakumar *et al* (27) as direct vPIC targets. In contrast to L genes, Latent and IE genes were mildly down-regulated.

**Fig.3.**
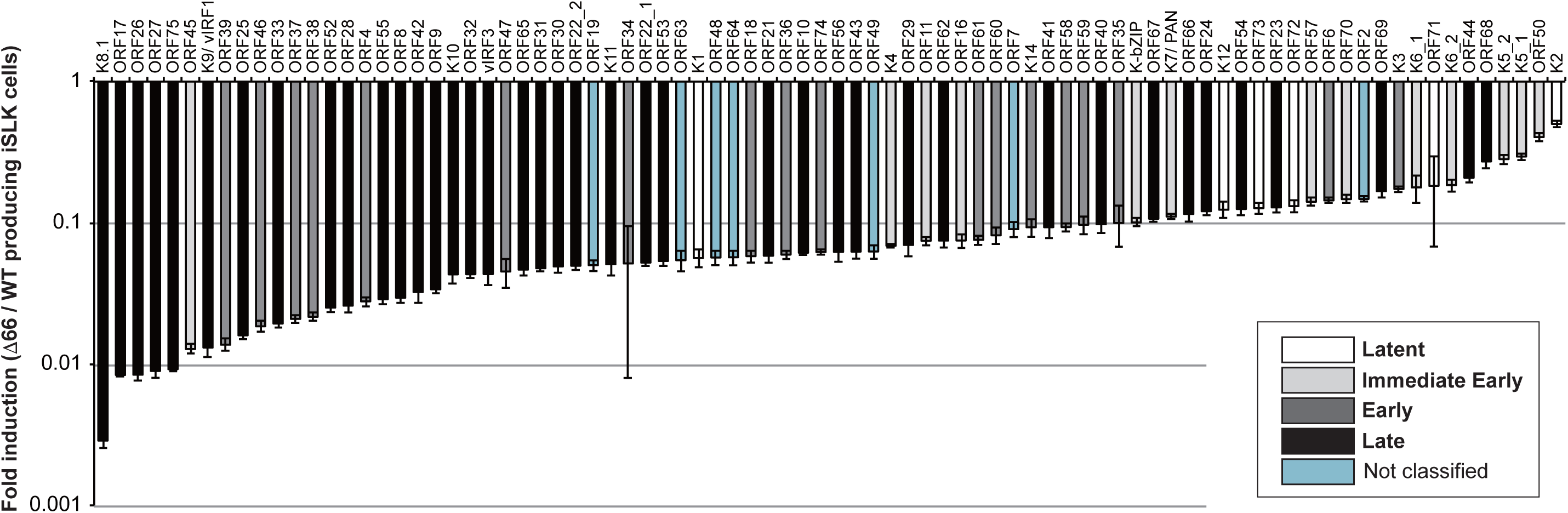
ORF66 is required for the late gene expression. iSLK-WT and iSLK-ΔORF66 cells were cultured for 72h in media with NaB and Dox to induce a lytic state. Total RNA was extracted from cells and subjected to RT-qPCR. The mRNA expression level of each viral gene was normalized by GAPDH expression, and columns indicated fold changes of iSLK-ΔORF66 transcripts compared with iSLK-WT transcripts. Classification of KSHV genes was performed according to Arias C. *et al.* (45). White, light gray, dark gray, black, and light blue columns indicate Latent, Immeidiate early, Early, Late, and Not classified genes, respectively. Expression levels were assessed using three independent samples, and error bars indicate ± standard deviations.

If ORF66 functions as a vPIC component, we hypothesized that it would be recruited to the “TATT”-box, which is known to be located approximately 30 bp upstream of the transcription start site (TSS) of L genes (27). To demonstrate this, we evaluated whether ORF66 interacts with L gene promoters by ChIP assay. iSLK-ΔORF66 and iVero-ΔORF66 cells stably expressing 3xFLAG-ORF66 were treated with or without NaB and Dox to induce lytic infection, and then cells were subjected to ChIP assay using anti-FLAG antibody (Fig. 4). As a result, immunoprecipitated ORF66 was found to be bound to the promoters of K8.1 (L gene) but not to the promoters of latent gene (ORF72), IE gene (ORF16) and E gene (ORF46/47). These results indicated that a protein complex bearing ORF66 may interact with L gene promoters via an interaction between ORF24 and ORF66.

**Fig.4.**
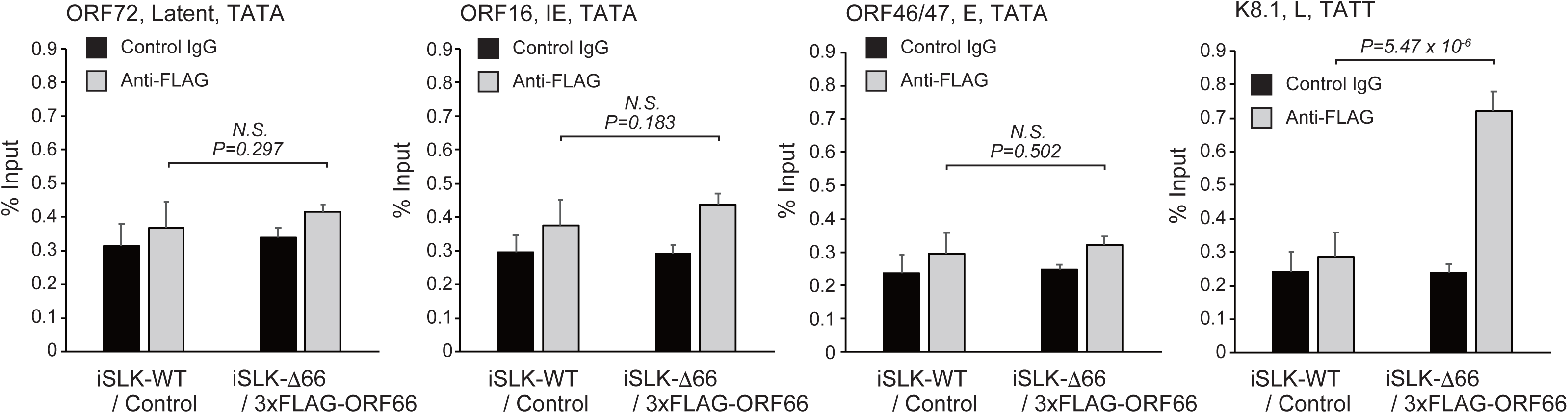
ORF66 associates with the transcriptional start site of Late genes. iSLK-WT/ Control and iSLK-ΔORF66/ 3xFLAG-ORF66 cells were treated with (or without) Dox and NaB for 72 hours and subjected to ChIP-qPCR. 3xFLAG-ORF66 protein was immunoprecipitated by anti-FLAG or control IgG antibody, and precipitates including chromatin and viral DNA were subjected to SYBR green real-time PCR for measuring the amount of promoter DNA of ORF72 (Latent gene), ORF16 (IE gene), ORF46/47 (E gene) or K8.1 (L gene). The levels of immunoprecipitated viral promoter were normalized to total input DNA.

### The interaction of ORF66-ORF34 and its function in viral production

ORF66 was reported to interact with other vPIC components such as ORF31, ORF18 and ORF34 by us and others (18, 20, 22). We also found that ORF34 operated as the vPIC hub, by bridging ORF24 and vPIC components including ORF66. Furthermore, these interactions are thought to be important for functions of vPIC in gene expression. To gain further insight into the interaction between ORF66 and ORF34, we identified the responsible region within ORF66 for binding with ORF34. We made ORF66 truncated mutants that each had approximately 60 amino acid deletions spanning from the N-terminus to C-terminus of ORF66 (Fig. 5a). The interaction between ORF66 mutants and ORF34 was assayed by pull-down experiments (Fig. 5b). 293T cells were co-transfected with 2xS-ORF66 mutants and 6xMyc-ORF34 plasmids, and 2xS-ORF66 were precipitated from cell extracts by S-protein-immobilized beads. As a result, ORF66 Δ1-Δ4 truncated mutants interacted with ORF34. However, no interaction with ORF66 Δ5-Δ7 truncated mutants was observed, meaning that the C-terminal region (241a.a.-429a.a.) of ORF66 is critical for ORF34 binding. Furthermore, we performed a *trans*-complementation assay using these truncation ORF66 mutants to assess how the loss of ORF34-ORF66 interaction affects virus production. ORF66 mutant plasmids were transfected into iSLK-ΔORF66 and iVero-ΔORF66, and recovery of virus production was measured. Wild type ORF66 expression significantly increased viral production in iSLK-ΔORF66 and iVero-ΔORF66 cells, while recovery of virus production was not detected by expressing any of the ORF66 truncated mutants (Fig. 5c and 5d). These results indicate that not only the ORF66 C-terminal domain (i.e., ORF34-binding region) but also the entire structure of ORF66 are indispensable for virus production.

**Fig.5.**
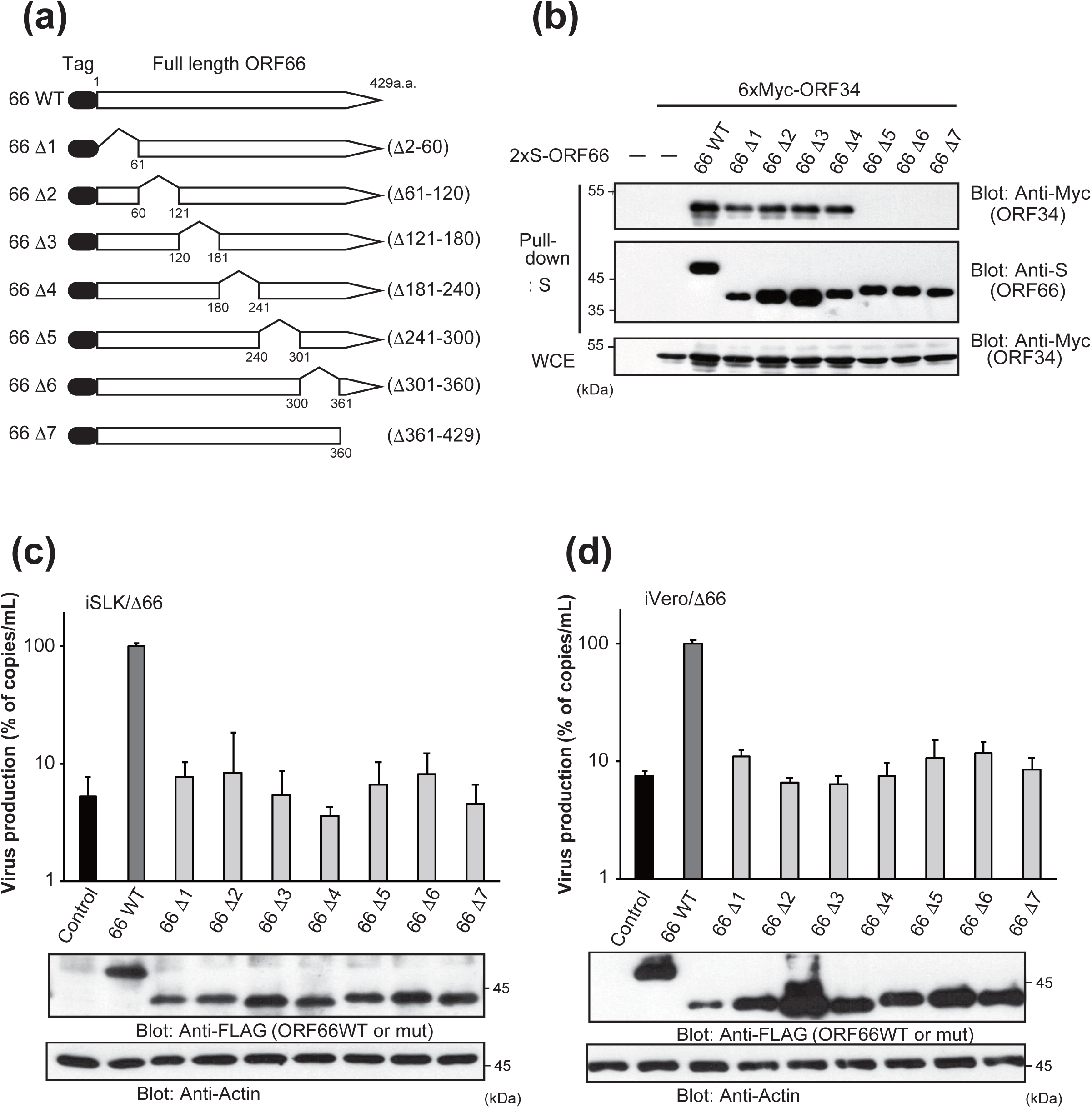
ORF66 physically interacts with ORF34 via its C-terminus regions and the entire structure of ORF66 is indispensable for virus production. (a) Schematic representation of tagged ORF34 deletion mutants used in the mapping experiments. The truncated amino acids are numbered next to its corresponding mutant. (b) 293T cells were co-transfected with expression plasmids of 2xS-ORF66 truncated mutant and 6xMyc-ORF34. Transfected cells were lysed, and cell lysates were subjected to pull-down assays using S-protein-immobilized beads that capture the 2xS-ORF66. Obtained precipitates including 2xS-ORF66 truncated mutants were probed with indicated antibodies to detect interactions. The (c) iSLK-ΔORF66 or (d) iVero-ΔORF66 cells were transfected with control, ORF66 or ORF66-truncated mutant plasmid. Transfected cells were stimulated for 3 days. Progeny KSHV was purified from harvested culture supernatant, and the KSHV genome was quantified by real-time PCR. Viral productivity was assessed using four independent samples, and error bars indicate standard deviations. Transfected virus producing cells were lysed and subjected to Western blotting to confirm ORF66 mutant expression.

An amino-acid sequence alignment of the ORF66 C-terminal domain of KSHV and other herpes virus homologs are depicted in Fig. 6a. Conserved amino acids are indicated with a gray background. To identify the key residues in the ORF66 C-termial region for interaction with ORF34, we constructed block alanine-scanning mutants (CR1mut to CR9mut), where several neighboring conserved amino-acids were substituted to alanine (Fig. 6a), except for proline to avoid disruption the whole protein structure. These mutants were subjected to pull-down assays to investigate the association of ORF34. Results showed that ORF66 CR2, 3, 4 and 6 mutants bind to ORF34, whereas ORF66 CR1, 5, 7, 8 and 9mut did not bind (Fig. 6b). Therefore, we focused on specific amino acid sequences of mutants losing the ability to bind to ORF34. In particular, CR1, 5, and 9mut have mutations in the C-X-X-C consensus sequence. On the other hand, CR7 contains a single cysteine residue, and CR8 contains three leucine residues.

**Fig.6.**
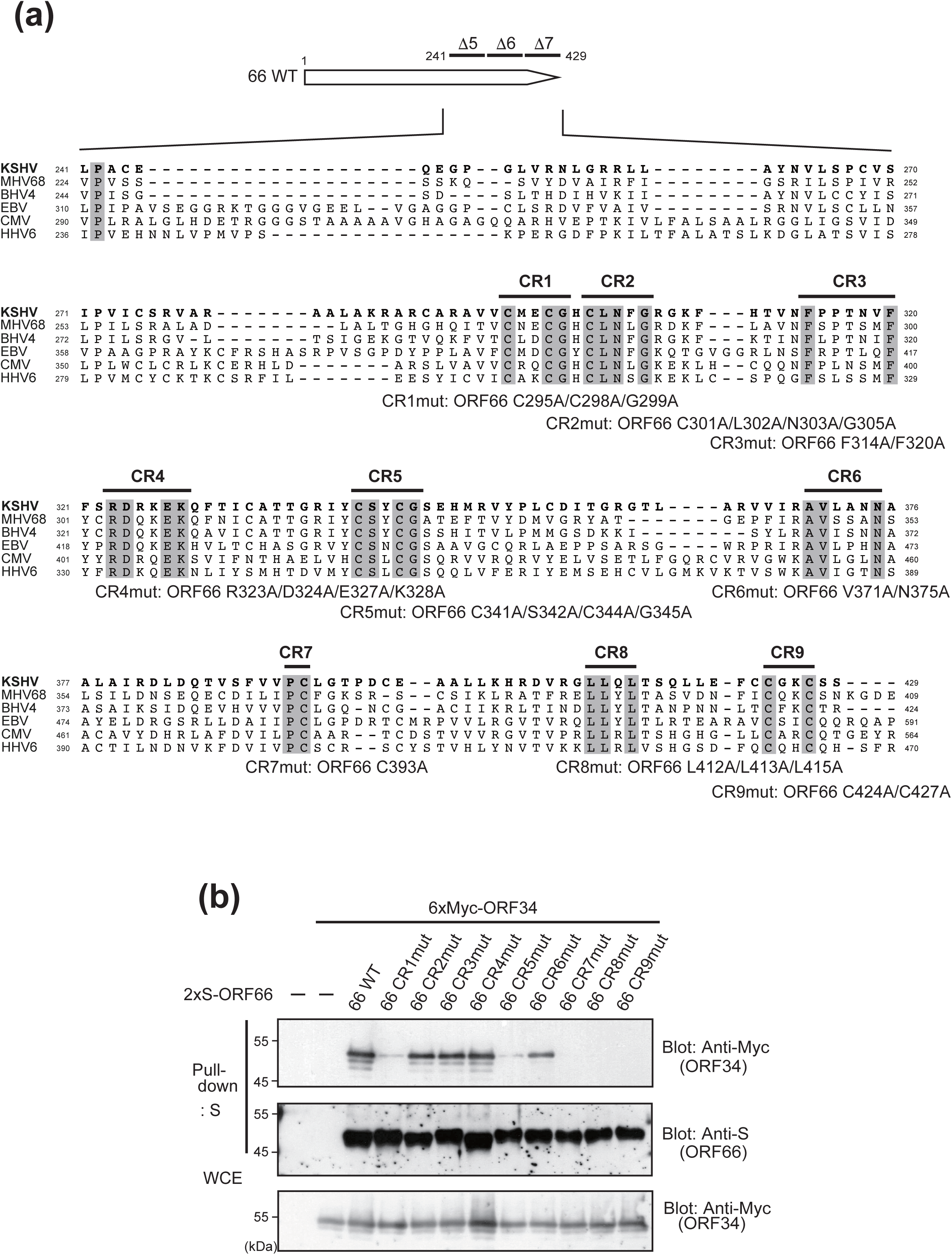
Several conserved residues of ORF66 are essential for physical association with ORF34. (a) Amino acid sequence alignment of C-terminus of ORF66 (241 a.a.-429 a.a.). Herpesvirus homolog amino acid sequences were translated from nucleotide sequences found in the NCBI database (KSHV ORF66 (JSC-1-BAC16; Accession number GQ994935), MHV68 ORF66 (strain WUMS; NC_001826), BHV4 ORF65 (strain V; JN133502), EBV BFRF2 (strain B95-8; V01555), CMV UL49 (strain Towne; FJ616285), HHV6 U33 (strain japan-a1; KY239023)). Raw data of alignment were obtained by using Clustal Omega (EMBL-EBI; https://www.ebi.ac.uk/Tools/msa/clustalo/). Completely conserved amino acids between homologs were indicated by a gray background. Based on this information, several conserved amino acids were split into blocks of alanine scanning ORF66 mutants (CR1mut; ORF66 C295A/C298A/G299A, CR2mut; ORF66 C301A/L302A/N303A/G305A, CR3mut; ORF66 F314A/F320A, CR4mut; ORF66 R323A/D324A/E327A/K328A, CR5mut; ORF66 C341A/S342A/C344A/G345A, CR6mut; ORF66 V371A/N375A, CR7mut; ORF66 C393A, CR8mut; ORF66 L412A/L413A/L415A, CR9mut; ORF66 C424A/C427A). (b) Block alanine scanning mutants were co-transfected with expression plasmids of 6xMyc-ORF34. Cell lysates were subjected to pull-down assays using S-protein immobilized beads.

To obtain farther insight into ORF34 binding via CR1 to CR9 of ORF66 in viral production, ORF66 alanine substitution mutants (CR1 to CR9) were subjected to *trans*-complementation assay using iSLK-ΔORF66 and iVero-ΔORF66 cells (Fig. 7a and 7b). Compared with ORF66 wild type expression, virus production in both iSLK-ΔORF66 and iVero-ΔORF66 cells could not be recovered by the expression of CR1, 5, 7, 8 and 9mut, which failed to interact with ORF34 (Fig. 6b). On the other hand, expression of CR2, 3, 4 and 6mut that maintained interaction with ORF34 showed varying levels of recovery, indicated by red columns (Fig. 7a-b). CR3 and 6mut showed almost the identical levels of recovery compared with ORF66 wild-type in iVero-ΔORF66 cells and partial recovery in iSLK-ΔORF66 cells. Recovery by the CR4mut was significant, however, lower than those of CR3 and 6mut. CR2mut had no effect. These data revealed that the binding between ORF66 and ORF34 via the conserved amino-acids of CR1, 5, 7, 8 and 9 regions in ORF66 is necessary, but not sufficient for virus replication.

**Fig.7.**
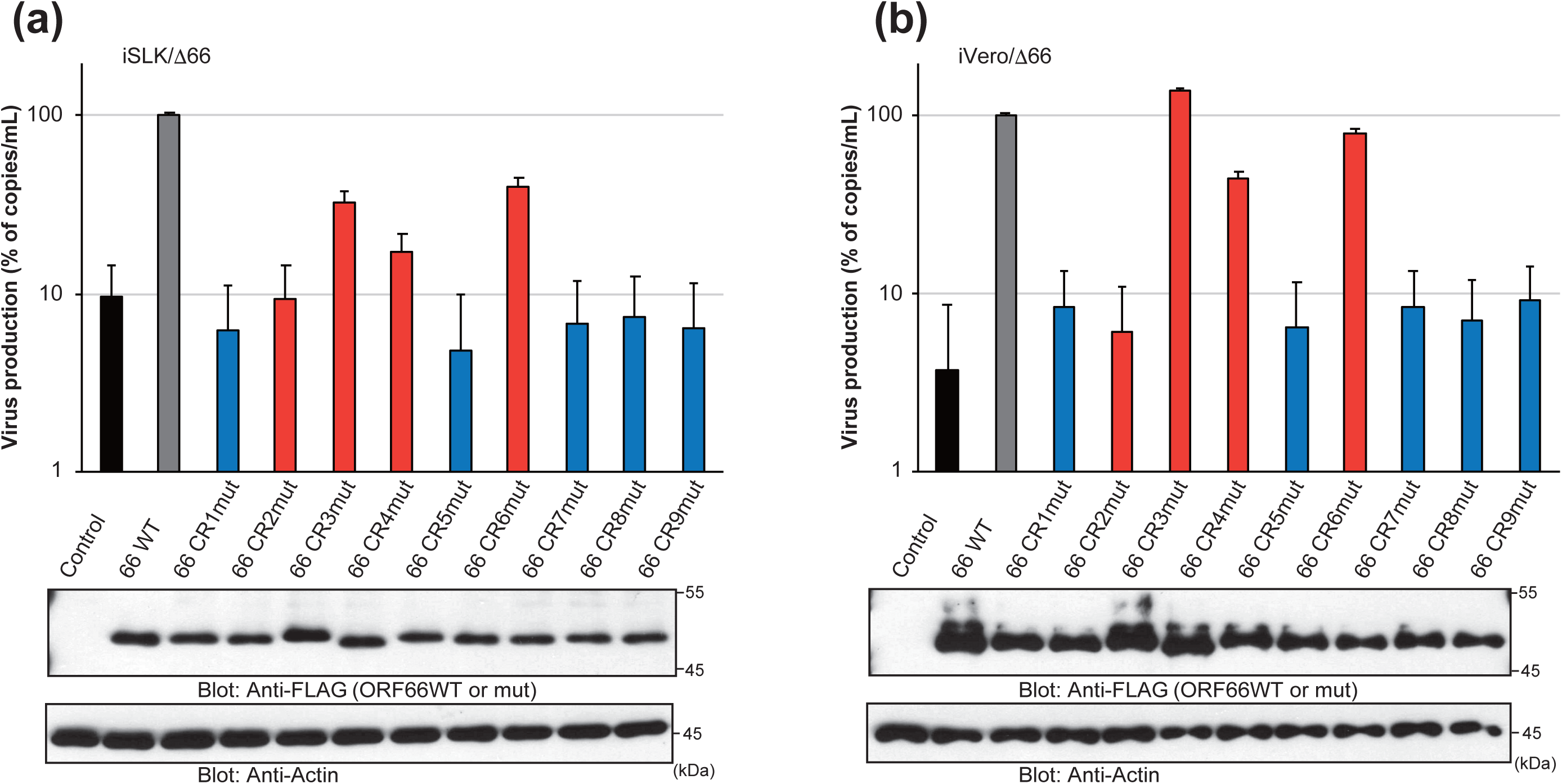
Association between ORF66 and ORF34 is necessary but not sufficient for virus production. The (a) iSLK-ΔORF66 or (b) iVero-ΔORF66 cells were transfected with control, ORF66 or ORF66 block alanine scanning mutant plasmid. Progeny KSHV was purified and the KSHV genome was quantified. Viral productivities were assessed using three independent samples, and error bars indicate standard deviations. The colour of each bar indicates the following: Black, control; Gray, ORF66 WT; Blue, ORF66 alanine scanning mutants not binding to ORF34; Red, ORF66 alanine scanning mutants binding to ORF34. Transfected virus producing cells were lysed and subjected to Western blotting analysis for the confirmation of ORF66 mutant expression.

### The C-X-X-C sequences and Zinc binding of ORF66 are important for association with ORF34

Based on our results of ORF66 alanine scanning mutants, we found that CR1, CR5 and CR9 regions of ORF66 contain a C-X-X-C sequence which is critical for ORF34 binding (Fig. 6a). Therefore, we generated expression plasmids with single amino acid mutants ORF66 C295A, C298A, C341A, C344A, C424A and C427A in which a cysteine in C-X-X-C was substituted with an alanine. Plasmids of ORF66 G299A, S342A, G345A and G345A mutants in which the conserved glycine or serine around the C-X-X-C sequence was substituted with alanine were also generated. The plasmids of 6xMyc-ORF34 and ORF66 alanine mutants were co-transfected into cells, and cell extracts were subjected to affinity purification using S-protein immobilized beads, followed by Western blotting. As a result, C295A, C298A, C341A, C344A, C424A and C427A ORF66 mutants failed to interact with ORF34 (Fig 8a, b and d). Furthermore, two leucine residues at 412 and 413 in CR8 of ORF66 were also important for the association with ORF34 (Fig 8c). These results suggest that a conserved leucine-repeat (412L and 413L) in CR8 and three conserved C-X-X-C sequences in CR1, 5 and 9 are needed for binding to ORF34, leading to the appropriate formation of vPIC. Because the C-X-X-C motif in proteins is often related to the binding to bivalent-cation, we speculate that C-X-X-C sequence contributes to form a higher order structure such as a zinc-finger domain. Next, to elucidate whether capturing zinc ion within ORF66 is necessary for binding to ORF34, we performed pull-down assays using zinc chelator TPEN and 2xS-ORF66-conjugated S-protein beads. The ORF66-conjugated beads were mixed with the cell extracts including overexpressed Myc tagged ORF34 in the presence of TPEN and were pull-down. The precipitates were subjected to western blotting with anti-Myc. As a result, TPEN decreased the interaction of ORF66 with ORF34, indicating that ORF66 may be a zinc-binding protein (Fig. 8e).

**Fig.8.**
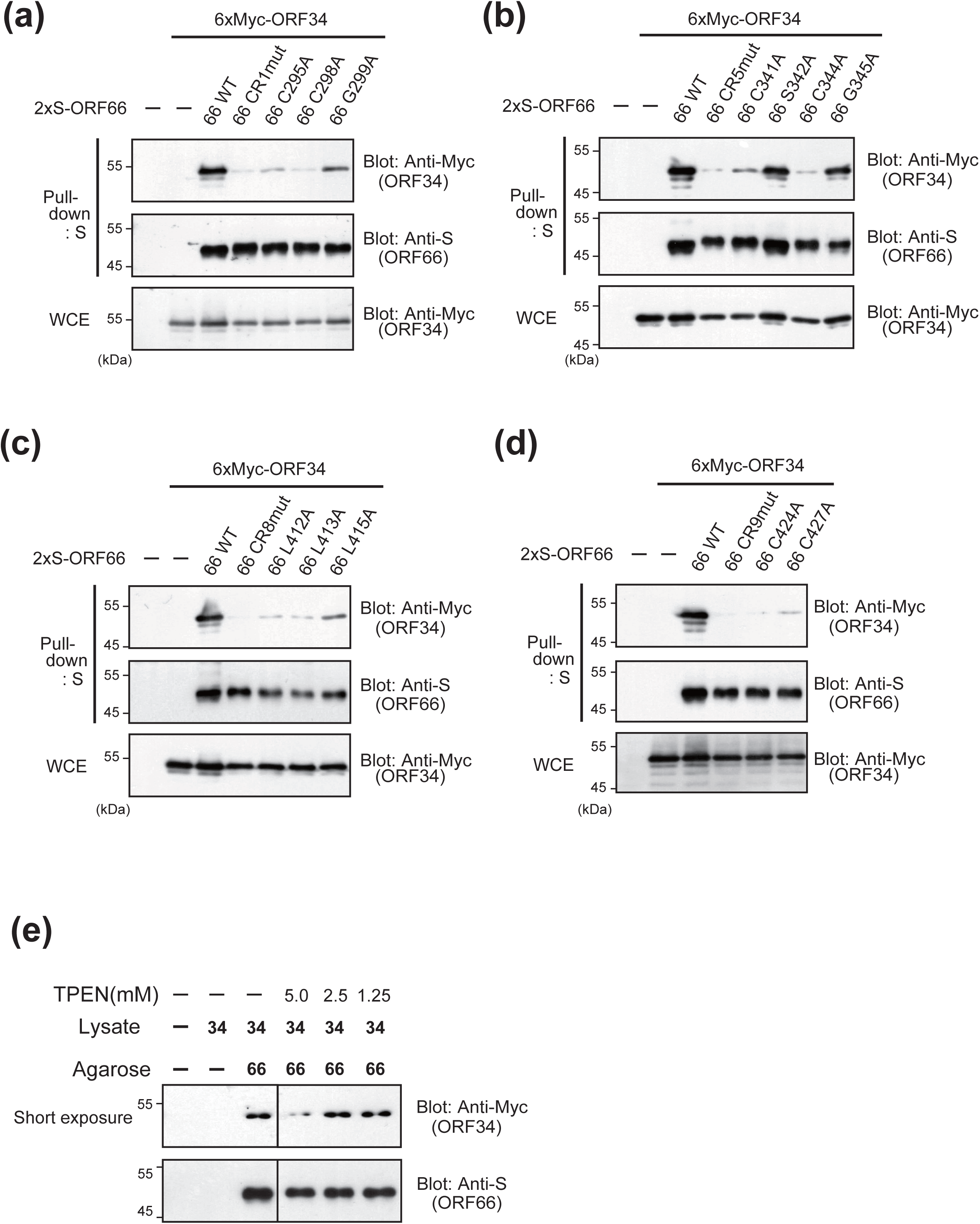
Identification of individual amino acids of ORF66 responsible for binding to ORF34. ORF66 wild-type, block alanine scanning mutants and single alanine scanning mutants were co-transfected with expression plasmids of 6xMyc-ORF34. Cell lysates were subjected to pull-down assays using S-protein immobilized beads. Blotting showed the association between ORF34 and (a) CR1 mutants (CR1mut; ORF66 C295A/C298A/G299A, ORF66 C295A, ORF66 C298A, ORF66 G299A, (b) CR5 mutants (CR5mut; ORF66 C341A/S342A/C344A/G345A, ORF66 C341A, ORF66 S342A, ORF66 C344A, ORF66 G345A), (c) CR8 mutants (CR8mut; ORF66 L412A/L413A/L415A, ORF66 L412A, ORF66 L413A, ORF66 L415A) and (d) CR9 mutants (CR9mut; ORF66 C424A/C427A, ORF66 C424A, ORF66 C427A). (e) Zinc ion chelation influences ORF66 binding abilities to ORF34. Pull-down assay using S-tagged ORF66 binding beads in the presence of Zinc chelator TPEN.

## Discussion

The C-terminus of ORF66 is highly conserved among herpesvirus homologs compared with the N-terminus (Fig.6a). We prepared alanine-scanning mutants of ORF66 for amino acid residues that are conserved among MHV, BHV, EBV, HCMV, and HHV-6. The conserved amino acid regions of CR1, 5, 7, 8 and 9 in the ORF66 C-terminal domain is necessary for the binding between ORF66 and ORF34, and virus production. In addition, three conserved C-X-X-C sequences in ORF66 CR1, 5 and 9 were needed for binding to ORF34 (Fig. 8). Our results indicate that ORF66 leads to appropriate vPIC formation by the interaction of ORF34 via C-X-X-C sequences in the C-terminal domain of ORF66. We speculate that the conserved C-X-X-C sequences could form a higher order structure such as a zinc-finger domain. Therefore, we approached the protein structure and functional prediction of ORF66 by a server-based helical protein structure simulation, I-TASSER (Iterative Threading ASSEmbly Refinement) (28–30) and generated a full-length homology model of ORF66 (Fig. 9). According to a meta-server approach to protein-ligand binding site prediction (COACH) (31, 32), four cysteine residues, C295, C298, C341, and C344 are predicted to associate with a zinc ion. Based on the location of each cysteine residue in the homology model, a pair of cysteines, C295/C298 in CR1/CR2 or C341/C344 in CR5, might chelate a single molecule of zinc, which correlates with our experimental observation using a zinc chelator, TPEN (Fig. 8). The zinc chelator TPEN inhibited association between ORF66 and ORF34. This result indicates that ORF66 binds zinc, which is important for its interaction with ORF34. We also performed homology searching of ORF66 by SWISS-model. Interestingly, 319-348 amino acids (including CR5 region) of ORF66 have low homology with the TFIIB zinc-ribbon domain of hyperthermophilic archaea *Pyrococcus furiosus*. As these zinc-associated cysteine residues were mostly located inside the protein structure, except for C341 (Fig. 9; middle panel), it is implies that CR1, CR2, and CR5 residues are involved in maintaining the functional structure formation of ORF66 via zinc interaction.

**Fig.9.**
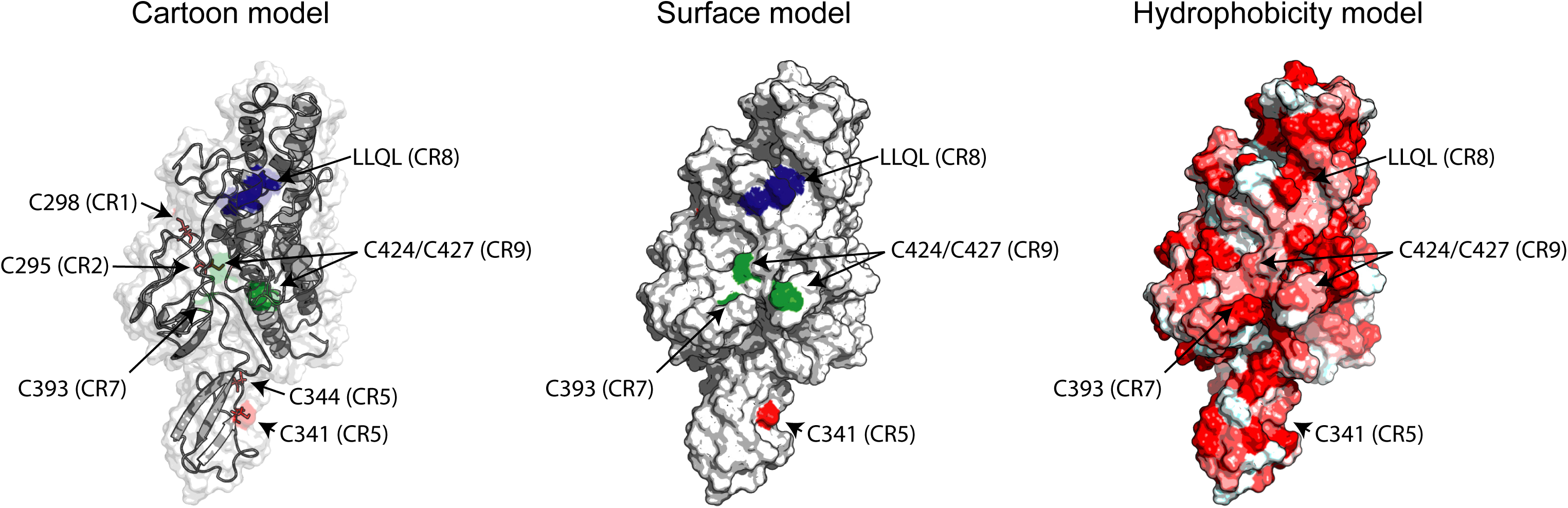
The protein structure and functional prediction of ORF66. The structural models of ORF66 simulated by I-TASSER, the server completed protein structural and functional prediction, are shown. The cartoon model indicates the position of responsible residues for interaction with ORF34 (left panel). The surface model represents the exposed residues on the protein surface (middle panel). The hydrophobicity of each residue is shown by a red color gradient followed by the Eisenberg hydrophobicity scale (right panel). The predicted Zinc ion-binding cysteins (C295 and C298, C341 and C344) are highlighted with brown. C344 in CR5 is highlighted in magenta. The LLQL in CR8 and C341 in CR5 are shown in cyan and green, respectively.

Two leucine residues (L412 and L413) in LLQL of CR8 are essential for binding between ORF66 and ORF34 (Figure 6b and 8c), suggesting that the hydrophobicity of a leucine-rich sequence is also indispensable for ORF66 to be functional. In fact, L412 and L413 residues in CR8 are predicted to be fully exposed on the protein surface, which represented a high degree of hydrophobicity by our structural model (Fig. 9). Two cytosine residues (C424 and C427) in CR9 were not predicted to be a zinc-binding domain by our protein-ligand binding prediction, however these residues in ORF66 were also responsible for ORF34 interaction (Figure 6). These residues are exposed on the protein surface (Figure 9; middle panel), and located at the hydrophobic surface including LLQL motif (Figure 9; right panel). Thus, the hydrophobic region surrounding two leucine residues in CR8 and cysteine residues in CR9 is expected to make up the binding surface and be directly involved in the hydrophobic interaction with ORF34.

We evaluated the viral replication in KSHV ΔORF66-producing iVero and iSLK cells, which were stably integrated with an ORF66-dificient KSHV BAC clone. ORF66 plays a critical role in virus production and the transcription of L genes. KSHV ORF24 binds to ORF34, RNA pol II and the TATT-box of the transcriptional start site (TSS) of the L gene (18, 20, 21). Because direct or indirect interaction of ORF34 with ORF 18, 30, 31 and 66 were indicated by co-immunoprecipitation and split luciferase experiments (18, 20, 21), ORF34 has been thought to function as a hub for interactions between ORF24 and other vPIC components. Therefore, these reports, in addition to our viral replication kinetics and viral gene expression data imply that ORF66 engages in L gene expression as a vPIC component. These results are in line with other herpesvirus homologs of KSHV ORF66. For instance, EBV BFRF2 is essential for virus production and contributes to vPIC formation (33). HCMV UL49 is also essential for replication in human foreskin fibroblasts (34, 35).

To gain a better understanding of ORF66 within the vPIC complex, we attempted to search for ORF34-binding regions within ORF66. Pull-down assays using truncated ORF66 mutants showed that the C-terminal region (from 241 a.a. to 429 a.a/the C-terminus end) of ORF66 was responsible for binding to ORF34 (Fig. 5b). And the truncated ORF66 mutants were subjected to a *trans*-complementation assay. However, all truncated ORF66 mutants did not rescue virus productions in both KSHV-ΔORF66 iVero and iSLK cell lines (Fig. 5c and 5d). Therefore, we speculate that the entire structure of ORF66 is necessary for virus production through vPIC formation. Considering the complex structure of human PIC components (TBP and GTFs) and RNAPII (36), vPIC is might be a crowded complex that consists of not only vPIC factors but also host proteins such as RNAPII, RNAPII binding proteins, and other unknown host. Lacking large regions of ORF66 may also influence the overal structure of the protein and affect interaction with its binding partners. Another possibility is that the N-terminal and center regions of ORF66 are related to interaction with other vPIC components or unknown host factors.

To evaluate the physiological function of conserved amino acids, block alanine-scanning mutants of ORF66 were subjected to a *trans*-complementation assay. The ORF66 mutants (CR1, 5, 7, 8 and 9mut) failed to associate with ORF34 (Fig. 6) and could not rescue virus production in KSHV-ΔORF66 cell lines (Fig. 7). In contrast, some ORF66 mutants (CR3 and 6mut), which associate with ORF34, showed full or partial rescue activities, while CR2mut did not. Presumably, the conserved sequence (CLNxG) in the CR2 region is related to binding to other vPIC associated factors. ORF66 rescued activity in iSLK/ Δ66 and iVero/ Δ66 cell lines, however, recovery rates were lower in iSLK/ Δ66. The differences in the recovery rates of both cell lines may be due to KSHV production potential and/ or characteristics of each cell line. Efficiency of KSHV production in iSLK is 100-fold higher than iVero. Furthermore, iVero is derived from non-human primates, African green monkey. Slight differences of mutated sites structure and/or species differences of host factors may influence the vPIC formation and accumulation of other host factors, resulting in differences in in the recovery rates. Altogether, association between ORF66 and ORF34 is necessary but not sufficient for virus production.

Our results show that the importance and molecular machinery of ORF66 in viral replication and L gene expression, as a vPIC component. Herpesvirus vPICs consist of viral factors as well as RNAPII and several host factors that are engaged in a complex. In MCMV, RNA helicase and cellular factors relating to splicing and translation interact with vPIC (37). Regulation of vPIC occurs through post-transcriptional modification of vPIC components, such as phosphorylation, which is known to contribute to physiological functions of vPIC in EBV (38). As the dynamics of vPIC in viral replication, vPIC target promoters on KSHV genome has more complexity than estimated. It inextricably linked to genome DNA replication (27). Our efforts to unveil vPIC machinery help to shed light on why β- and γ-herpesviruses have incorporated vPIC machinery into its genome for survival.

## Material and Methods

### Plasmids

pCI-neo-3xFLAG-ORF66, pCI-neo-3xFLAG-ORF66, pCI-neo-6xMyc-ORF34 expression plasmids were previously described (20). Truncation and alanine mutant ORF66 coding fragments were obtained by PCR or overlap-extension PCR from ORF66 expression plasmid using primer sets noted in Table 1, and were cloned into pCI-neo-2xS, and pCI-neo-3xFLAG vectors respectively. For pCI-blast plasmid construction, Blastcidin resistant gene (blaR) coding fragments were obtained by PCR from pLKO.1-blast (Addgene plasmid # 26655, a kind gift from Dr. Keith Mostov. (39)) using primer sets as noted in Table 1, and were replaced Neo^R^ in pCI-neo mammalian expression vector (Promega, WI, USA). pCI-neo-3xFLAG and pCI-neo-3xFLAG-ORF66 were digested with NheI and NotI sites, and the protein coding fragments were inserted into MCS of pCI-blast to construct of pCI-blast-3xFLAG and pCI-blast-3xFLAG-ORF66.

**Table 1:**
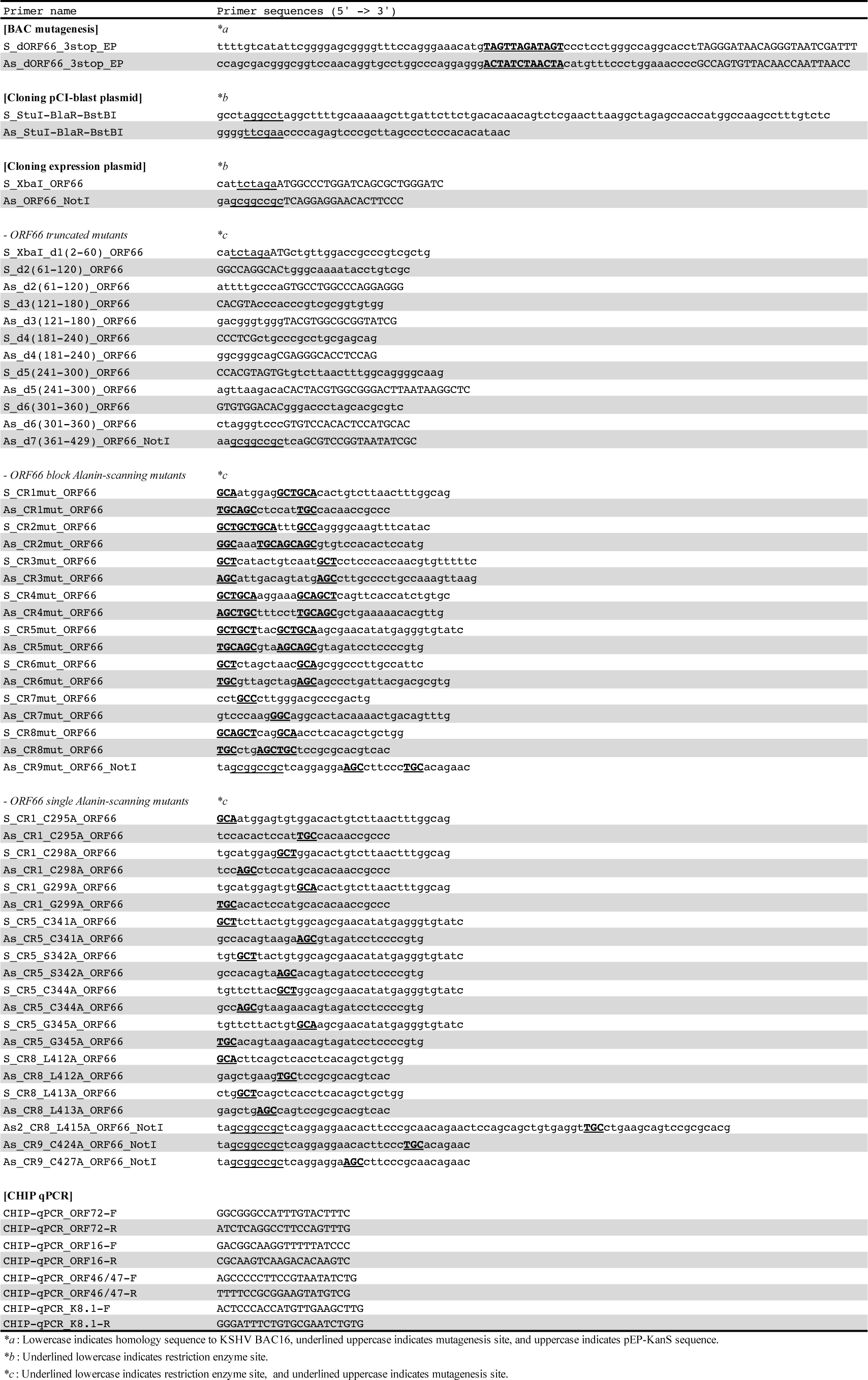
Primers for construction of expression plasmids

### Mutagenesis of KSHV BAC16

KSHV BAC16 was a kind gift from Jae U. Jung and mutagenesis of KSHV BAC16 was performed according to previous publications (24, 25). The primers for mutagenesis sequences are noted in Table 1. Insertion and deletion of kanamycin resistance cassettes (KanR) in each mutant were analyzed by digestion of EcoRV and agarose-gel electrophoresis. Mutated sites of each BAC clone were confirmed by Sanger sequencing.

### Establishment of doxycycline-inducible recombinant KSHV-expressing cells and stably ORF66-expressing cells

For maintenance, iSLK cells were cultured in growth medium containing 1 μg/mL of puromycin (InvivoGen, CA, USA) and 0.25 mg/mL of G418 (Nacalai Tesque, Kyoto, Japan). iVero cells (20) were cultured in a growth medium containing 2.5 μg/mL of puromycin. KSHV BAC16 wild-type (WT-BAC16) and mutant (ΔORF66-BAC16) were transfected to iSLK and iVero cells. iSLK and iVero cells were transfected by a calcium phosphate and a lipofection method, respectivly. Transfected cells were selected under 1000 μg/mL of hygromycin B (Wako, Osaka, Japan) and 2.5 μg/mL of puromycin to establish doxycycline-inducible recombinant KSHV producing cell lines (iSLK-WT, iSLK-ΔORF66, iVero-WT, iVero-ΔORF66).

To establish stable ORF66-expressing cells, pCI-blast-3xFLAG-ORF66 and empty vector (pCI-blast-3xFLAG) were transfected into iSLK-WT or iSLK-ΔORF66 cells, and transfected cells were selected in 10 μg/mL of Blastcidin S (InvivoGen) and maintained in 7.5 μg/mL of Blastcidin S. Thus, stable cell lines, iSLK-WT/pCI-blast-3xFLAG, iSLK-ΔORF66/pCI-blast-3xFLAG and iSLK-ΔORF66/pCI-blast-3xFLAG-ORF66, were established. To establish iVero-WT/pCI-neo-3xFLAG, iVero-ΔORF66/ pCI-neo-3xFLAG, iVero-ΔORF66/ pCI-neo-3xFLAG-ORF66 stable cell lines, iSLK harboring each KSHV BAC clone was selected and maintained in 1.5 mg/mL of G418.

### Measurement of virus production and viral DNA replication

For quantification of virus production, KSHV virions in culture supernatant were quantified as previously described (20, 40, 41). Briefly, iSLK and iVero cells (iSLK-WT, iSLK-ΔORF66, iVero-WT, or iVero-ΔORF66) were treated with Sodium Butyrate (NaB) and doxycycline (iSLK; NaB 0.75 mM/ Dox 4 μg/mL, iVero; NaB 1.5 mM/ Dox 8 μg/mL) for 72 hours to induce to lytic replication and production of recombinant KSHV, and culture supernatants were harvested. Culture supernatants (220 μL) were treated with DNase I (NEB, MA, USA) to obtain only enveloped and encapsidated viral genomes. Viral DNA was purified and extracted from 200 μL of DNase I-treated culture supernatant using the QIAamp DNA blood mini kit (QIAGEN, CA, USA). To quantify viral DNA copies, SYBR green real-time PCR was performed using KSHV-encoded ORF11 specific primers.

For measurement of KSHV genome replication, each KSHV producing cell line was treated with doxycycline and NaB for 48 hours to induce lytic replication, and harvested. Total cellular DNA containing the KSHV genome were purified and extracted from washed cells using the QIAamp DNA blood mini kit (QIAGEN). Cellular KSHV genome copies were determined by SYBR green real-time PCR and normalized to total DNA.

### Recovery of exogenous gene in BAC harboring cells

The iSLK cells were transfected with pCI-neo-3xFLAG as a control plasmid, and 3xFLAG-tagged ORF66 full length, truncated mutant, alanine mutant plasmids using Screenfect A plus (Wako, Tokyo, JAPAN) according to the manufacturer’s instructions and simultaneously stimulated with NaB 0.75 mM / Dox 4 μg/mL containing medium. The iVero cells were transfected using PEI-MAX MW40000 (Polysciences, Inc., Warrington, PA, USA)(42). After 1 day, transfected iVero cells were stimulated with NaB 0.5 mM / Dox 8 μg/mL containing medium. After three days of stimulation, viral supernatant was harvested and KSHV genome was evaluated by real-time PCR.

### RT real-time PCR (RT-qPCR) array

mRNA was extracted from iSLK cells treated with Dox and NaB using FastGene RNA Premium Kit (Nippon Genetics Co. Ltd., Tokyo, Japan). cDNA was synthesized by ReverTra Ace RT-qPCR kit (TOYOBO, Osaka, Japan) and subjected to SYBR green real-time PCR. Gene expression was analyzed by qPCR using specific primers designed by Fakhari and Dittmer (26). Relative KSHV mRNA expression levels were determined by GAPDH expression and ΔΔCt methods.

### Chromatin Immuno-precipitation (ChIP) assay

ChIP assay was performed as described previously (43) with slight modifications. Briefly, iSLK-ΔWT/ Control and iSLK-ΔORF66/ 3xFLAG-ORF66 cells were treated with or without 4 μg/mL of Dox and 0.75 mM NaB for 72 hours. Formaldehyde-fixed cells were lysed by farnham lysis buffer (5 mM PIPES pH 8.0 / 85 mM KCl / 0.5% NP-40) and the nuclear pellet was collected. The pellet was lysed in SDS lysis buffer and sonicated. The supernatant containing DNA was diluted by CHIP dilution buffer and then subjected to immunoprecipitation with anti-FLAG (DDDDK-tag) monoclonal antibody (FLA-1; MBL, Nagoya, Japan) or mouse control IgG (Santa-Cruz). Immunoprecipitates containing chromatin and viral DNA were subjected to SYBR green real-time PCR for measuring the amount of promoter DNA of each gene. The amount of immunoprecipitated viral DNA was normalized to 1% of input DNA. The sequences of qPCR primer sets for each transcription start site of ORFs are noted in Table 1.

### Pull-down assay, Western blot, and antibodies

Western blots were performed as described previously (20). For pull-down assays, transfected 293T cells (RCB2202; RIKEN Bio Resource Center, Tsukuba, Japan) were lysed by HNTG buffer (20 mM HEPES (pH 7.9), 0.18 M NaCl, 0.1% NP-40, 0.1 mM EDTA, 10% Glycerol) with protease inhibitors and sonicated. The cell extracts were subjected to affinity purification using S-protein immobilized beads (Novagen, MA, USA), and purified proteins (containing 2xS-tagged ORF66 or mutants) were subjected to western blotting.

For Zinc chelator TPEN (N,N,N’,N’-Tetrakis (2-pyridylmethyl) ethylenediamine; TCI, Tokyo, Japan) treatment, 3xFLAG-taggged ORF66 overexpressed in 293T cells, was purified with S-protein immobilized beads in the presence of each dose of TPEN or vehicle (ethanol) for 2 hours. The beads were mixed with cell lysate of 6xMyc-ORF66 overexpressed in 293T in the presence of each dose of TPEN or vehicle (ethanol) for 2 hours. The beads were washed 4 times and subjected to western blotting.

Anti-Myc (9E10; Santa-Cruz, CA, USA), Anti-S-tag pAb (MBL, Nagoya, Japan), Anti-FLAG (DDDDK-tag) (FLA-1; MBL), Anti-Actin (AC-15; Santa-Cruz) were used as the primary antibodies. HRP linked anti-mouse IgG antibody (GE Healthcare UK Ltd., Buckinghamshire, UK) or HRP linked anti-rabbit IgG antibody (GE healthcare UK Ltd.) was used as the secondary antibody. Anti-FLAG-HRP (M2; Sigma-aldrich, MO, USA) was also used.

### Homology modeling

The template structure for ORF66 was initially identified by collecting high-scoring structural templates from Local Meta-Threading Sever to generate a 3D structural model. The protein-ligand predictions were then derived by threading the 3D models through a protein function database, BioLiP. All procedures were automatically processed on I-TASSER server program provided by the Zhang lab (URL: https://zhanglab.ccmb.med.umich.edu/I-TASSER/) (28–32). The visualization of the homology model was performed by molecular visualization open-source software, PyMOL. The hydrophobicity in each amino acid was determined by the Eisenberg hydrophobicity scale (https://web.expasy.org/protscale/pscale/Hphob.Eisenberg.html) (44) and visualized by running a color_h.py script on PyMOL.

### Statistics

The two-tailed student’s t-test was used to indicate the differences between the groups. *P* values are shown in each figure.

## Acknowledgements

The BAC16, KSHV BAC clone, was a kind gift from Dr. Kevin Brulois and Dr. Jae U Jung (U.S.C., US). We thank Dr. Gregory A. Smith (Northwestern Univ., US) for the E. coli strain GS1783, and Dr. Nikolaus Osterrieder (Cornell Univ., US) for the plasmid pEP-KanS. We thank Dr. Peter Gee for scientific advice and critical proofreading of the manuscript. This work was supported in-part by a Grant-in-Aid for Scientific Research (C) (18K06642), Young Scientists (B) (16K18925) and Young Scientists (18K14910) from the Ministry of Education, Culture, Sports, Science and Technology of Japan.

